# Enzalutamide-induced PTH1R-mediated TGFBR2 decrease in osteoblasts contributes to resistance in prostate cancer bone metastases

**DOI:** 10.1101/829044

**Authors:** Shang Su, Jingchen Cao, Xiangqi Meng, Ruihua Liu, Alexandra Vander Ark, Erica Woodford, Reian Zhang, Isabelle Stiver, Xiaotun Zhang, Zachary B. Madaj, Megan J. Bowman, Yingying Wu, H. Eric Xu, Bin Chen, Haiquan Yu, Xiaohong Li

## Abstract

Over 80% of prostate cancer (PCa) patients in the United States die with bone metastases. Second-line hormonal therapies, such as enzalutamide, improve overall survival in about 50% of patients with bone metastases, but almost all responsive patients eventually develop enzalutamide resistance. Our study showed that although enzalutamide significantly inhibited the tumor growth of subcutaneously or orthotopically grafted PCa C4-2B cells, it had no effect on the bone lesion development when C4-2B tumors were grafted in the bone, suggesting a crucial role of the microenvironment in enzalutamide resistance in PCa bone metastasis. We found that enzalutamide significantly decreased the amount of the TGFBR2 (TGF-β type II receptor) in osteoblasts, both *in vitro* and in patient samples. The osteoblast-specific knockout of *Tgfbr2* significantly induced bone metastasis. We showed that the enzalutamide-induced TGFBR2 decrease in osteoblasts was mediated by increased PTH1R (parathyroid hormone/parathyroid hormone-related peptide receptor), which resulted in TGFBR2 degradation, and that blocking PTH1R rescued the TGFBR2 decrease. Furthermore, we found that PTH1R up-regulation by enzalutamide was correlated with increased *Pth1r* promoter occupancy by transcription factor NR2F1. Our findings highlight a potential enzalutamide-resistance mechanism through TGFBR2 decrease in osteoblasts, thus suggesting future PTH1R-blocking approaches to overcome enzalutamide resistance in PCa bone metastasis.

## Introduction

Prostate epithelial cells depend on androgen for growth, and because androgen signaling stimulates most prostate cancer (PCa) cell growth, androgen deprivation therapy (ADT) is the first-line treatment for prostate cancer (1–3). Enzalutamide, a second-generation small-molecule inhibitor of the androgen receptor (AR), has been approved for patients who develop castration-resistant PCa bone metastases (mCRPC) after ADT failure (4–6). However, resistance developed within an average of 6 months. Therefore, it is an urgent need to study the enzalutamide resistance in PCa bone metastasis (7, 8).

Cancer cells are genetically instable, and many enzalutamide-induced resistance mechanisms in cancer cells have been identified both in patients and in laboratory models, including intrinsic resistant mechanisms in AR negative adenocarcinoma or neuroendocrine cells, the ability of AR positive cells to transdifferentiate into AR negative neuroendocrine cells, or the possibility that AR negative cells were selected by enzalutamide-induced killing of AR positive cells (4, 9–11). Altogether, the heterogeneity and plasticity of cancer cells make targeting only cancer cells unlikely to achieve the goal of overcoming resistance. Therefore, we aim to study the non-malignant cells in the tumor microenvironment, i.e. bone cells, and to identify the enzalutamide-regulated events that contribute to bone metastases (4). By identifying the intrinsic changes or downstream molecular events that result from enzalutamide treatment in the bone microenvironment, specifically in the niche-forming cells such as osteoblasts, we may develop a novel therapeutic strategy to overcome enzalutamide resistance in PCa bone metastasis and thus improve therapeutic outcomes.

Consistent with previous studies (12), we showed that enzalutamide inhibited the tumor growth of C4-2B PCa cells in subcutaneous or orthotopic xenografts. However, enzalutamide treatment had no effect on the development of bone lesions induced by C4-2B cells that were injected into the mouse tibiae. These data suggested a crucial role of the microenvironment in enzalutamide resistance in PCa bone metastasis. We then found that enzalutamide significantly reduced the expression of the TGF-β type II receptor (TGFBR2) in cultured osteoblasts, as well as in osteoblasts from PCa patients who had been treated with second generation AR inhibitors such as enzalutamide. Previously, we showed that osteoblast-specific knockout of the TGF-β type II receptor gene (*Tgfbr2*) stimulated bone metastasis of AR-negative PCa cells (12). In this study, we found that specific knockout of *Tgfbr2* in osteoblasts also promoted the bone metastasis of C4-2B cells, which are AR-positive cells and castration-resistant (13). These data suggested a potential drug resistance mechanism using the decrease of TGFBR2 in osteoblasts by enzalutamide. Focusing the molecular mechanism of this regulation, we demonstrated that the events responsible for the enzalutamide-induced TGFBR2 decrease in osteoblasts include the increased expression of the transcription factor NR2F1 (Nuclear Receptor Subfamily 2 Group F Member 1) followed by the NR2F1-activated increase of PTH1R (parathyroid hormone/parathyroid hormone-related peptide receptor), which induced TGFBR2 degradation. Blocking PTH1R rescued the decrease of TGFBR2 by enzalutamide. These data suggest a possible strategy of using PTH1R blockade to overcome enzalutamide resistance in PCa bone metastasis.

## Results and Discussion

The majority (80%) of PCa bone metastasis samples express AR (14). Therefore, to determine whether the effect of enzalutamide on PCa depends on the microenvironment, we used AR-positive prostate cancer C4-2B cells, which are castration-resistant but still responsive to androgen and further AR inhibition such as enzalutamide (15). One million C4-2B cells were injected subcutaneously into the flank (Figure 1A), intratibially (Figure 1B), or orthotopically into the prostate **(Supplemental Figure 1)** of 5-to 6-week-old male SCID mice. Enzalutamide (20 mg/kg body weight) or vehicle was administered daily through oral gavage for consecutive 2 to 3 weeks, starting from day 15 (the subcutaneous and orthotopical models) or day 1 (the intratibial model) post injection. The dosing was chosen based on previous publication (16). We conducted all the statistical analyses in these three experiments using a linear mixed-effect model and found that the C4-2B cells responded differently to enzalutamide in the different microenvironments. As expected, enzalutamide treatment significantly inhibited both subcutaneous (Figure 1A, *p* = 0.001, 8.3% slower rate of increase in lesion area in the treated group, 95% CI = 3.4 – 13%) and orthotopic C4-2B tumor growth (**Supplemental Figure 1**, *p* = 0.023). However, enzalutamide treatment had no effects on bone lesion development in the mice intratibially injected with C4-2B (Figure 1B, *p* = 0.793, 4.3% higher rate of increase in lesion area in the treated group, 95% CI = −42.7 – 24.8%). In other words, PCa cells in the bone microenvironment were resistant to enzalutamide, suggesting that the bone microenvironment likely caused this enzalutamide resistance.

**Figure 1.**
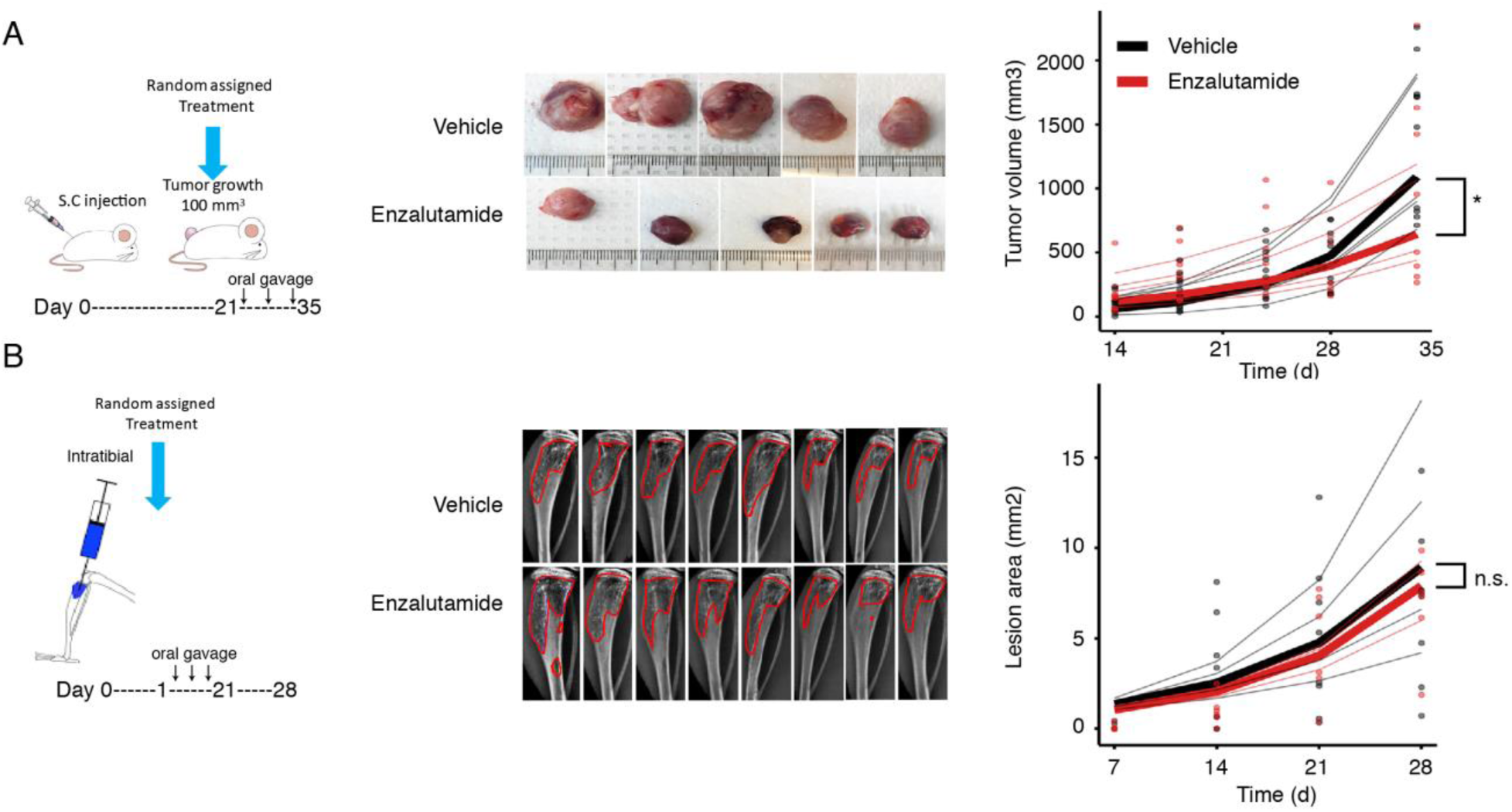
Enzalutamide treatment was effective in subcutaneous models of prostate cancer but not in intratibial models. **A)** Subcutaneous tumor volume progression and representative pictures of individual tumors are shown. The bold lines are the mean values of the individual curves of each color. These parallel experiments were repeated twice with the same results. Enzalutamide dissolved in vehicle (15% DMSO + 85% PEG300), or vehicle alone was administered by oral gavage. Two weeks of treatments were started on day 21 when the tumor volume reached 100 mm^3^. Tumor size was measured twice per week. *p* = 0.001 using a linear mixed-effect model (*n* ≥ 5 xenografts). **B)** Bone lesions were imaged by X-ray weekly, and lesion areas were measured using Metamorph software. The lesion areas measured are outlined. Treatments were started on day 2 after cancer cell injections. *p* = 0.793 using a linear mixed-effect model (*n* ≥ 8 tibiae). If not specified, * indicates *p* <0.05 and n.s. indicates non-significant (*p* ≥ 0.05).

We then focused our mechanistic studies on osteoblasts because metastatic PCa cells attach to them when colonizing in the bone (17, 18). It has been reported that the normal cell culture medium with 10% fetal bovine serum (FBS), which contains hormones such as androgen, mimics the microenvironment with very low level of androgen seen in the clinical situation under ADT (19). Therefore, we cultured murine MC3T3-E1 osteoblasts under these conditions to determine the effect of enzalutamide on osteoblasts. As mentioned above, we recently identified the role of osteoblast-specific *Tgfbr2* loss in promoting the bone metastasis of AR-negative PC-3 and DU145 cells (12). We then checked the expression of TGFBR2 in enzalutamide-treated osteoblasts and found a decreased amount of TGFBR2 at 24 to 72 h post enzalutamide treatment and at 20 μM and 40 μM doses. Enzalutamide at 40 μM also decreased TGFBR2 at 8 h and 12 h post enzalutamide treatment. No change in TGFBR2 was observed at 6 h of treatment (Figure 2A). To validate the specificity of enzalutamide effects, we knocked down the *Ar* gene in MC3T3-E1 osteoblasts and observed a consistent decrease of TGFBR2 (Figure 2B), confirming the on-target effect of enzalutamide. The doses of enzalutamide were chosen based on previously reported Cmax of enzalutamide (35 μM) (20).

**Figure 2.**
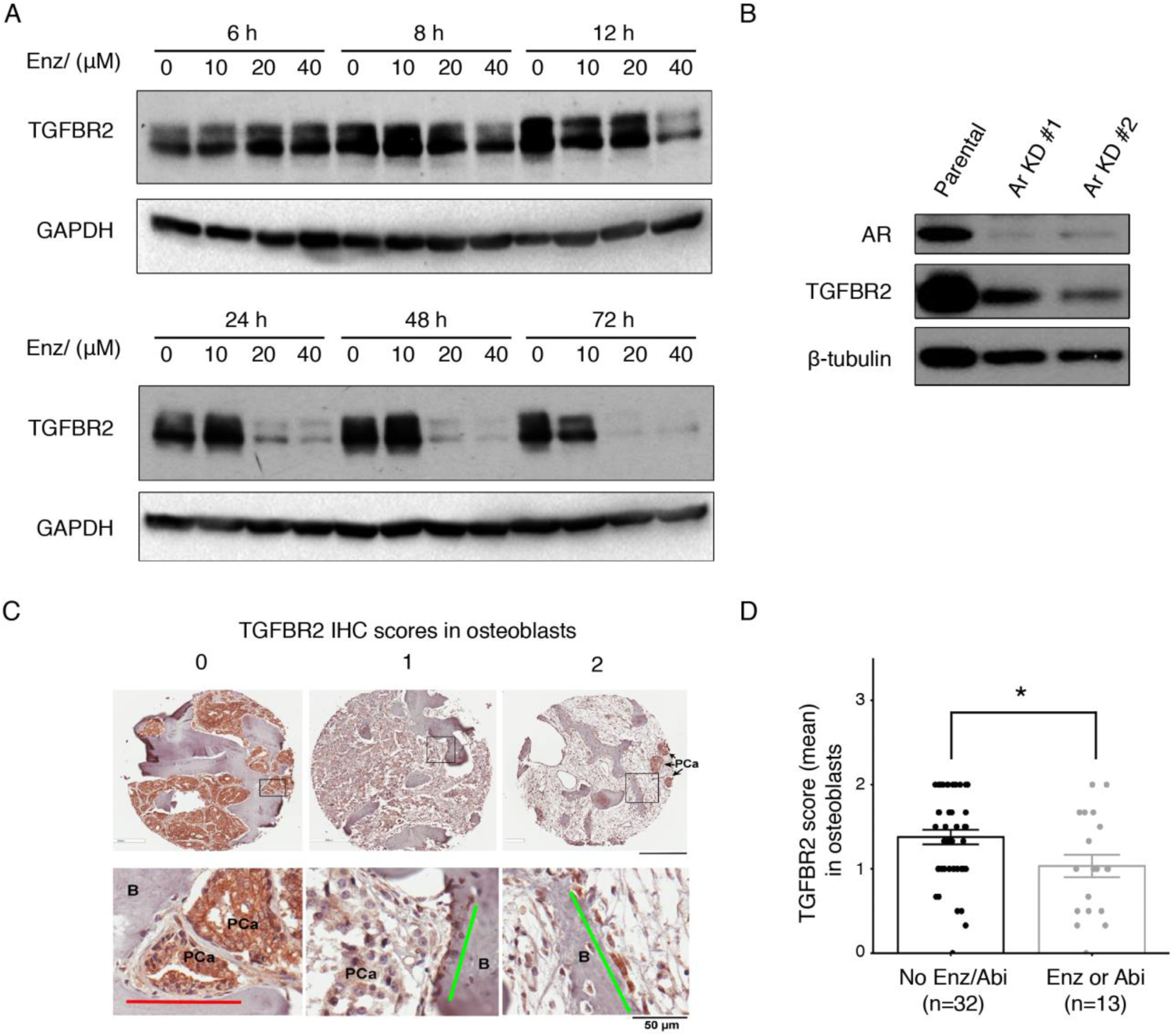
Enzalutamide decreased TGFBR2 in osteoblasts. **A)** Time-and dose-dependent effects in MC3T3 cells. 0 = vehicle (DMSO). **B)** AR-dependent effect of TGFBR2. The mixed cell populations (but not selected single-cell clones) of *Ar* knockdown (KD) were achieved using the CRISPR-Case9 system. Patental, cells without guide RNA manipulation. Experiments were repeated three times with the same results. **C)** Representative IHC images of TGFBR2 in osteoblasts. Lines in zoomed lower panel highlight osteoblasts. PCa, prostate cancer cells; B, bone Scale bar, upper panel 300 μm, lower panel 50 μm. **D)** Mean IHC scores for TGFBR2 in osteoblasts in different patient groups. Enz, enzalutamide. Abi, abiraterone. *p* = 0.03 using a linear mixed-effect model (n indicates patient number).

To determine whether the *Tgfbr2* KO in osteoblasts had a promoting effect on bone metastasis of AR-positive PCa, we injected C4-2B cells into the tibiae of the previously established mouse model *Tgfbr2^Col1CreERT^* KO, in which *Tgfbr2* can be specifically knocked out in osteoblasts upon induction (12). C4-2B cells induced mixed (osteoblastic and osteolytic) lesions (**Supplemental Figure 2A**) (21). We found 21 out of 24 injected mouse tibiae developed C4-2B cell-induced bone lesions in the *Tgfbr2*^*Col1CreERT*^ KO group, whereas 7 out of 18 injected mouse tibiae developed bone lesions in the *Tgfbr2*^*FloxE2*^ (control) group (**Supplemental Figure 2B**). A two-sample proportions test suggested these two groups significantly (*p* < 0.001) differed in their proportions of the incidence of bone lesions. As for the bone lesion area analysis, we performed the Wilcoxon rank sum test with continuity correction and applied the Bonferroni-Holm multiple test correction. We found the knockout group had significantly greater (*p* = 0.0414) bone lesion area (median = 3.4996) than the control group (median = 0) at week 5 post injection, although the lesion areas in the two groups were not significantly different (*p* = 0.4261) when we excluded those tibiae without bone lesions (area = 0) (**Supplemental Figure 2C, D**). A similar pattern was observed when the lesion areas were examined at week 5.5 post injection (**Supplemental Figure 2C, D**).

To gain clinical insights into the TGFBR2 expression in PCa bone metastases, we performed an IHC screening on TGFBR2 levels in a set of mCRPC patient tissue microarrays (UWTMA79) supported by the Prostate Cancer Biorepository Network (PCBN). Statistical analysis using a mixed linear-effect model showed the expression scores of TGFBR2 in osteoblasts were significantly lower (*p* = 0.03) in patients who had received second-line AR signaling blockers (n = 13, either enzalutamide or abiraterone, an androgen biosynthesis inhibitor) than those who did not receive enzalutamide or abiraterone (n = 32) (Figure 2C, D) (22). No difference of TGFBR2 expression in other cell types (such as tumor cells, stromal/fibroblasts, or osteoclasts) was observed in these patients (**Supplemental Table 1**). No correlations of TGFBR2 expression in osteoblasts were found for any other treatment group (such as bisphosphonate medicine Zometa, or antifungal ketoconazole) (**Supplemental Table 1**). The data suggested a potential drug resistance in PCa bone metastasis from the enzalutamide-induced TGFBR2 decrease in osteoblasts.

The decrease of TGFBR2 by enzalutamide in MC3T3-E1 cells was not observed until 8 h of treatment (see Figure 2A), suggesting an indirect effect. One previous study has shown that parathyroid hormone (PTH) activates PTH1R to recruit TGFBR2, which phosphorylates PTH1R and which together undergo endocytosis for protein degradation (23). Bone metastatic PCa cells are known to secrete factors such as PTHrP (parathyroid hormone-related protein), which could also bind and activate PTH1R but would give a distinct conformation relative to PTH-activated PTH1R (24, 25), suggesting that PTHrP could induce TGFBR2 degradation by activation of PTH1R.

To observe changes of endogenous proteins, we treated MC3T3-E1 cells with the protein synthesis inhibitor cycloheximide together with PTHrP. We found that both TGFBR2 and PTH1R were decreased to an undetectable level after 12 h of PTHrP treatment (Figure 3A). TGF-β stimulation results in the phosphorylation of Smad2 (p-Smad2), the activation of downstream signaling, and the degradation of TGFBR2 through ubiquitination(26); so TGF-β treatment was used as a positive control for p-Smad2 up-regulation and TGFBR2 down-regulation(Figure 3A). These data suggested that PTHrP, through activation of PTH1R, induced the down-regulation of TGFBR2 and its downstream signaling.

**Figure 3.**
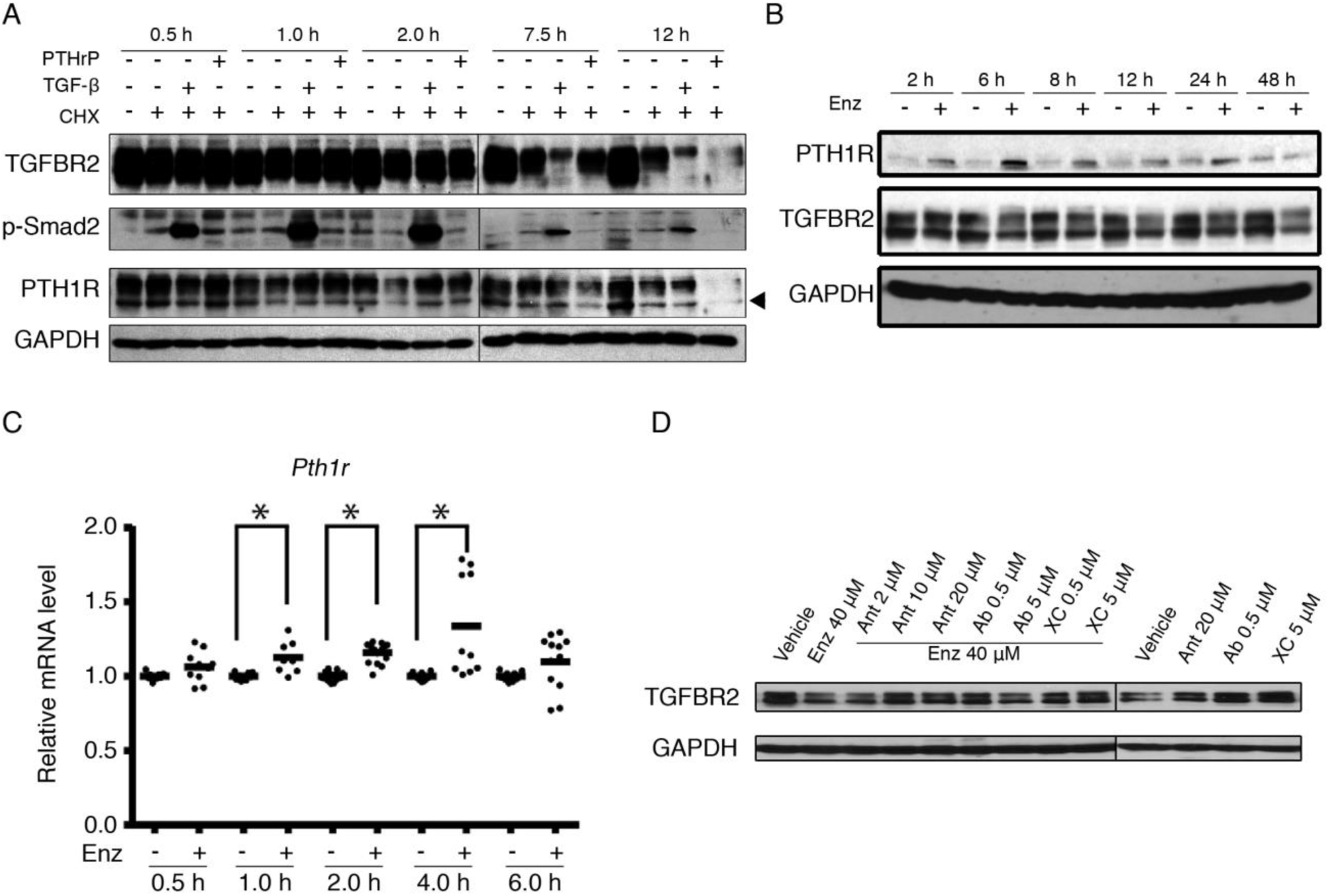
Enzalutamide-induced TGFBR2 down-regulation was mediated by PTH1R. **A)** TGFBR2 down-regulation with PTH1R. MC3T3-E1 cells were treated with 10 μg/mL cycloheximide (CHX), 1 ng/mL TGF-β, or 100 nM PTHrP at the indicated times. Total protein extractions from cells were used for western blotting. Arrowhead indicates major band of PTH1R. **B & C)** Enzalutamide increased PTH1R at the protein **(B)** and mRNA **(C)** levels prior to the decrease of TGFBR2. MC3T3-E1 cells were treated with vehicle (-, DMSO) or enzalutamide (enz, 40 μM). Total proteins were extracted from cells for western blotting. The scatter plot in **C** displayed mean and individual values generated from multiple biological and technical replicates. * indicates *p* < 0.05 using a linear mixed-effect model (n≥8 from duplicate wells per biological samples). **D)** Blockade of PTH1R rescued the TGFBR2 loss upon enzalutamide treatment. MC3T3 cells were treated with vehicle, DMSO; Ant, the peptide antagonist for PTH1R; Ab, the neutralizing antibody of PTHrP; or XC, the small-molecule inhibitor of PTH1R for 24 h before immunoblotting. All the western blotting experiments were repeated three times with the same results.

We then conducted a time-course treatment of enzalutamide on MC3T3-E1 cells and found that prior to the decrease of TGFBR2, enzalutamide treatment increased PTH1R (Figure 3B). PTH1R proteins were increased upon enzalutamide treatment at multiple time points from 2 to 24 h but were not affected at 48 h. The most significant increase in PTH1R was observed at 6 h, when there was no change of TGFBR2. TGFBR2 down-regulation was consistently observed at and after 24 h of treatment. These data suggested that enzalutamide might directly increase PTH1R, which subsequently result in the decrease of TGFBR2. We further observed a significant increase in *Pth1r* mRNA at early time points (1 h, 2 h, and 4 h; *p* <0.05 using a line mixed-effect model test) upon enzalutamide treatment (Figure 3C), suggesting that PTH1R up-regulation is one of the early effects of enzalutamide on osteoblasts. When we treated C4-2B PCa cells stably expressing TGFBR2 (termed as C4-2B/TGFBR2) or RAW264.7 osteoclasts with enzalutamide, however, PTH1R did not increase, although TGFBR2 did decrease at certain time points, suggesting that the mechanism by which enzalutamide-induced PTH1R-mediated TGFBR2 decrease only exist in osteoblasts (**Supplemental Figure 3**). This is important because TGFBR2 loss in osteoblasts stimulates bone metastasis but that loss in myeloid lineage cells inhibits bone metastasis(12). Functionally, blocking PTH1R rescued the TGFBR2 expression reduced by enzalutamide, either with a small-molecule inhibitor of PTH1R, a neutralizing antibody against PTH1R, or a peptide antagonist of PTHrP (Figure 3D). These data collectively demonstrated that PTH1R is a key player in the enzalutamide-induced TGFBR2 decrease.

Next, we aimed to determine the mechanism by which enzalutamide increased PTH1R in osteoblasts. As PTH1R was up-regulated at both the transcriptional and protein levels (see Figure 3B, C), the most likely explanation would be the enhanced transcriptional activation of *Pth1r* by transcription factor(s). We adopted a text-mining approach using MetaCore to search for candidates that are linked with both AR and PTH1R, and the top hit we got was Nuclear Receptor Subfamily 2 Group F Member 1 (NR2F1) (Figure 4A). NR2F1 was negatively associated with AR signaling and could activate the expression of *Pth1r* (27, 28). We observed a significant increase (1 h, 2 h, and 4 h; *p* <0.05 using a line mixed-effect model test) of *Nr2f1* mRNA in osteoblasts treated with enzalutamide (Figure 4B) and an increase of NR2F1 proteins (Figure 4C). We did not observe an increase of NR2F1 in RAW264.7 or C4-2B/TGFBR2 cells (**Supplemental Figure 3**). We downloaded the binding element information of NR2F1 (5-A/G-A/G-GGTCA-3) from the JASPAR database and searched the element across the upstream regulatory sequences of mouse *Pth1r* (Figure 4D) (29). There are at least three major transcript variants of mouse *Pth1r* (labeled as V1, V2, and V3, driven by at least three different promoters, as shown in Figure 4D); Promoter of V1 (containing Exons E1, E2, E4 and those follows) is considered absent or very weak in the bone while those of V2 (containing E3, E4, and those follows) and V3 (containing E4’, E4 and those follows) are active in the bone (30). We found multiple NR2F1 binding elements at the promoters of V2 and V3. We conducted ChIP-qPCR (primer amplicons termed as p1, for V2, and p2, for V3, as shown in Figure 4D) to determine the occupancy of NR2F1 in the *Pth1r* promoter region. We found that NR2F1 bound to both designed promoter regions, and the occupancy was enhanced by enzalutamide, indicated by the elevated fold enrichment (Figure 4D). These data suggested that enzalutamide induced PTH1R expression by enhancing the *Pth1r* promoter occupancy of NR2F1.

**Figure 4.**
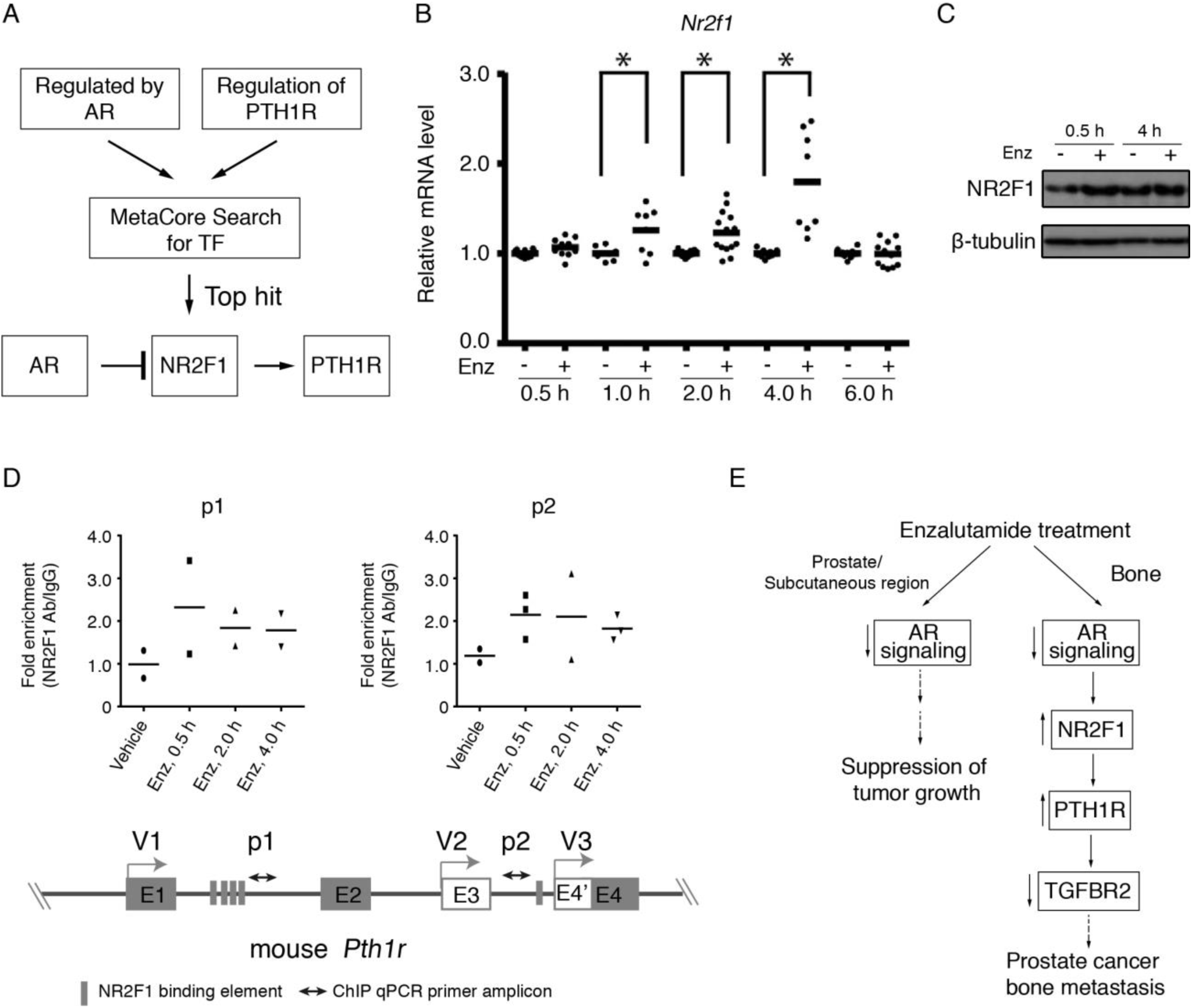
Enzalutamide induced NR2F1 expression and recruitment on the Pth1r promoter. **A)** NR2F1 was identified as a potential transcription factor responsible for PTH1R transcriptional activation upon AR signaling blockade**. B &C)** Enzalutamide up-regulated NR2F1 at both the mRNA **(B)** and protein levels **(C)** in osteoblasts. MC3T3 cells were treated with vehicle (-, DMSO) or enzalutamide (enz, 40 μM) for the indicated hours before harvest for immunoblotting or RNA extraction. The scatter plot in **B** displayed mean and individual values generated from multiple biological and technical replicates. * indicates *p* < 0.05 using a linear mixed-effect model (n≥8 same as above). **D)** Enzalutamide increased the occupancy of NR2F1 at the *Pth1r* promoter. NR2F1 binding elements appear frequently across *Pth1r* promoter(s) of different *Pth1r* transcript variants (V1, V2, and V3). ChIP assay with different targeting primers (amplicons termed as p1and p2) all showed an increase in NR2F1 occupancy on *Pth1r* regulatory regions upon enzalutamide treatment. Scatter plots displayed the mean and individual values from biological replicates. E1~E4 indicate exons in different variants. Note that the *Pth1r* genomic region plotted here was not proportional to actual length. **E)** Proposed mechanism of enzalutamide resistance in bone metastasis. In a responsive microenvironment, such as the prostate and subcutaneous region, enzalutamide blocks AR and suppress PCa tumor growth. In bone, enzalutamide blocks AR and removes the suppression of AR on NR2F1 expression. Elevated NR2F1 is then recruited onto the *Pth1r* promoter, which stimulates the expression of PTH1R. The PTH1R proteins induce the degradation of TGFBR2 in osteoblasts to promote prostate cancer bone metastasis.

Our study highlights the importance of understanding the role of the tumor microenvironment in PCa bone metastasis and drug resistance by demonstrating a signaling regulatory axis of enzalutamide specific in osteoblasts. Distinct from the enzalutamide-induced AR signaling blockade and tumor growth suppression in the prostate and subcutaneous region, enzalutamide blockade of AR increased the *Pth1r* promoter occupancy of NR2F1, stimulating the expression of PTH1R, which in turn led to the degradation of TGFBR2 in the bone microenvironment (Figure 4E). It usually takes several weeks to generate enzalutamide resistant cancer cell clones *in vitro*, while the enzalutamide induced PTH1R-mediated TGFBR2 degradation in osteoblasts were observed within 2 days. Our data thus suggests promising approaches to overcome enzalutamide resistance by blocking PTH1R.

## Methods

Experimental procedures are provided in **Supplemental Methods.**

*Study approval*. All experiments using SCID, *Tgfbr2*^*FloxE2*^ and *Tgfbr2*^*Col1CreERT*^ knockout (KO) mice complied with federal and institutional guidelines, and the research was approved by the Van Andel Institute (VAI) Institutional Animal Care and Use Committee (IACUC) under protocol numbers 19-01-002 and 10-01-001.

## Author contributions

XL conceived and directed the project. XL, SS, JC and YW wrote the manuscript. XL, SS, JC, and XM designed the experiments and analyzed the data with help from RL, AVA, EW, RZ, and IS. XZ scored the cell-specific TGFBR2 expression in tissue microarray. ZM and YW performed the statistical analyses. MB conducted the MetaCore analysis. EX, BC, and HY provided resources.

## Acknowledgements

We thank Drs. Bart Williams, Patrick Grohar, Tao Yang, Matt Steensma, and their lab members for intriguing discussion and technical help, and David Nadziejka for technical editing. This work was supported by NCI grant R01CA230744 and a grant from Van Andel Institute (50310A) to XL. The prostate cancer bone metastasis tissue microarray was purchased from the Prostate Cancer Biorepository Network (funded by W81XWH-14-2-0182, W81XWH-14-2-0183, W81XWH-14-2-0185, W81XWH-14-2-0186, and W81XWH-15-2-0062).

