## Supplemental Data for "Enzalutamide-induced PTH1R-mediated TGFBR2 decrease in osteoblasts contributes to resistance in prostate cancer bone metastases"

Contents:

- **Full methods**
- **Supplemental Table 1 (provided as a separate excel file）**
- **Supplemental Table 2**
- **Supplemental Figures 1-3**
- **References**

**Animals, cells, proteins, and drugs**

The genetically engineered mouse models, *Tgfbr2^FloxE2^* and *Tgfbr2^Col1CreERT^* knockout (KO), have been characterized and described previously (1). Briefly, the mice were generated by crossing collagen 1α2 promoter-driven CreERT (*Col1CreERT*) mice and *Tgfbr2^FloxE2^* mice. To activate the Cre recombinase, the *Cre*^–^ and *Cre^+^* littermates were injected with tamoxifen (10 mg/kg body weight) for five consecutive days. Between the 5^th^ and 7^th^ day after the last tamoxifen injection, the resulting *Tgfbr2^FloxE2^* (control) and *Tgfbr2^Col1CreERT^* KO mice were confirmed by genotyping and immunohistochemical analysis of the harvested tibiae. These mice were crossed to immunodeficient mice so that they could grow human tumors. The bones of the *Tgfbr2^Col1CreERT^* KO mice were normal, as analyzed by X-ray (no bone lesions or fractures), microCT (for bone mineral density, cortical and trabecular bones, cross-sectional bone thickness and parameter), and histomorphometry (for the number of osteoclasts and osteoblasts) (1). In this study, the mice were crossed to the NSG-SCID mouse background, so C4-2B PCa cells could grow in the bones. The research using these mice was approved by the Van Andel Institute (VAI) Institutional Animal Care and Use Committee (IACUC) under protocol numbers 19-01-002 and 10-01-001.

Mouse osteoblast MC3T3-E1 cells and human osteoblast hFOB1.19 cells were from Dr. Bart Williams’ lab of VAI, mouse macrophage RAW264.7 (used for osteoclasts) cells were from Dr. Cindy Miranti’s lab at Arizona State University, and human PCa C4-2B cells were obtained from Dr. Leland Chang at Cedars Sinai. C4-2B/TGFBR2 cells were generated using an expression construct that was kindly provided by Dr. Jindan Yu at Northwest University. All cells were grown in a humidified incubator with 5% CO_2_ at 37 °C, except for hFOB1.19 cells which were grown at 34 °C. The cells were periodically checked for mycoplasma contamination, and only mycoplasma-free cells were used for experiments.

Enzalutamide was purchased from Selleckchem and Sigma-Aldrich. Abiraterone was purchased from Selleckchem. XC039 or XC, the PTH1R small-molecule inhibitor, was provided by Dr. Eric Xu (Van Andel Institute). These drugs were all dissolved in DMSO at various concentrations for *in vitro* studies. For *in vivo* studies, enzalutamide was dissolved in vehicle, 15% DMSO + 85% PEG300. (Asn^10^, Leu^11^, D-Trp^12^) PTHrP(7–34) amide, a peptide antagonist for PTH1R, and neutralizing antibody of PTHrP were purchased from Bachem. All the details for the special reagents in this study can be found in Supplemental Data.

**Experimental mouse models**

NSG-SCID male mice, 6-8 weeks old, were purchased from the vivarium of VAI for use in this study. For subcutaneous xenograft models, 1 million C4-2B cells were suspended in 100 μL of Matrigel and subcutaneously injected into the flank of NSG-SCID mice. Tumors were measured twice per week using digital calipers. Tumor volume was calculated using the formula V = (*a*^2^ × *b*)/2, *a* and *b* indicating the minimal and maximal diameter, respectively. When the volume of subcutaneous tumors reached 100 mm^3^, mice were randomized into a treatment group (enzalutamide at 20 mg/kg body weight once per day through oral gavage) or a vehicle group (given the same volume of the vehicle, 15% DMSO + 85% PEG300, once per day via oral gavage). For orthotopic xenograft models, 1 million C4-2B cells were suspended in 10 μL of PBS and injected into the anterior lobe of the mouse prostate. Tumors were monitored once per week by bioluminescent imaging. Enzalutamide treatments started when the bioluminescent signals of orthotopic tumors were comparable to the signals from 100-mm^3^ subcutaneous tumors.

For intratibial injection, 1 million C4-2B cells in 10 μL of PBS were injected into the mouse tibiae as described previously (1). The injected mice were then randomized and started on drug treatment the next day. Bone lesions and tumor growth were monitored once per week by radiographic imaging using a Faxitron X-ray machine and bioluminescent imaging using an AMI-1000, respectively. The bone lesion areas and regions of interest (ROIs) were measured using MetaMorph (Molecular Devices Inc.).

Cell preparations, injections, imaging, tumor measurements, and bone lesion analyses were performed blinded.

**Western blotting**

Cells were lysed with RIPA lysis buffer containing Roche’s Complete Proteinase Inhibitor Cocktail Tablets. The total protein was quantified using the BCA protein quantitative method according to the manufacturer’s protocol. Approximately 10-40 μg protein per sample was loaded and electrophoresed through 10% or 12% SDS polyacrylamide gels and transferred onto nitrocellulose membranes. The membranes were then incubated overnight at 4 °C with primary antibodies for TGFBR2 (1:1000, Santa Cruz # sc-17792 or sc-400), p-Smad2 (1:1000, Cell Signaling # 3108), PTH1R (1:500, Abcam # 180762; 1:1000, Biolegend # 906401), NR2F1 (1:1000, Cell Signaling # 6364) or AR (1:500, Santa Cruz # sc-816). GAPDH (1:10000, Cell Signaling # 5174; 1:2000, Bimake #A5028), β-actin (1:5000, Sigma # A5441), or β-tubulin (1:5000, Sigma # T4026) was used as the loading control. HRP-conjugated secondary antibodies goat-anti-mouse # 31430 and goat-anti-rabbit # 31460, diluted 1:5000-10,000) and enhanced chemiluminescence (ECL) HRP substrate (# 34076) were purchased from ThermoFisher Scientific.

**CRISPR/Cas9-based gene editing of *Ar* in MC3T3-E1 cells**

Five different single guide RNAs (sgRNA) for mouse *Ar* were designed and synthesized in complementary DNA oligos. Annealed oligos were then inserted into digested pLentiCRISPRv2. The resulting five plasmids were confirmed by sequencing, then packaged into lentivirus and infected into MC3T3 osteoblasts. The infected cells were then selected with puromycin (2.5 μg/mL) for 7 days to get positive stable cell populations. Cells were harvested and western blots were performed to confirm the gene knockdown and TGFBR2 regulation. Results for two of the five MC3T3 populations derived from those independent gRNAs were shown in Figure 2B. The primer sequences for generating the gRNA oligos were shown in **Supplemental Table 2**.

**Tissue microarray, histology, and immunohistochemistry**

The PCa metastasis tissue microarray (TMA) UWTMA79 was obtained from the University of Washington. This TMA contained samples from 45 cases of bone and visceral metastasis from rapid autopsy (62 visceral and 79 bone metastasis). The slides were subjected to immunohistochemical analysis of TGFBR2, as described previously. The stained slides were digitally captured and quantified independently by the pathologist using a score of 0 to 3 on various cell types for intensities of negative, moderate, and strong. The scores were submitted to the Bioinformatics and Biostatistics Core for discovery analyses of correlations among the cell-specific expression of TGFBR2 and any of the clinical parameters.

Subcutaneous or orthotopic C4-2B tumors, as well as C4-2B injected tibiae, were harvested at the endpoints and fixed in 10% neutral-buffered formalin for 2 d. After fixation, the tibiae were processed to decalcification in 14% EDTA for 5-6 days. All the steps were performed at 4 °C. The tissues were then paraffin-embedded. Paraffin-embedded sections (5 μm) were deparaffinized and stained with hematoxylin and eosin (H&E). For immunohistochemistry, paraffin-embedded sections were subjected to standard peroxidase-based immunohistochemistry procedures. The TGFBR2 antibody (Santa Cruz # sc-17792) was used with 1:50 dilution.

**RNA extraction and qRT-PCR**

Total RNA was isolated using TRIzol (Invitrogen) and subsequently purified using the RNeasy kit (Qiagen). Complementary DNA was synthesized using a SuperScript VILO cDNA Synthesis Kit (Invitrogen), and qRT-PCR was performed using SYBR Select Master Mix (Bio-Rad) with an ABI 7500 machine (Applied Biosystems). Cycle threshold values were determined and normalized to the loading control, *Gapdh*, for each experiment. Fold changes for experimental groups were calculated using the ΔΔC_T_ method. The primer sequences are listed in **Supplemental Table 2**.

**Chromatin immunoprecipitation (ChIP) analysis**

ChIP analysis was conducted with a Simple ChIP Kit from Cell Signaling Technology (#9003), following the manual. Briefly, 4 million adherent MC3T3 cells treated with either vehicle (DMSO) or enzalutamide were rinsed with ice-cold PBS once to remove the residual culture medium and then were fixed with 1% formaldehyde in PBS at room temperature for 15 min on a gentle horizontal shaker. Glycine was then introduced into the solution for 5 min to terminate the fixation. The liquid was aspirated, and the cells were rinsed twice with cold PBS. The cells were scraped with cold PBS and centrifuge at 500 × *g* and 4 °C for 2 min to pellet the cells. Then the cells were lysed, digested with MNase, and sonicated to release the chromatin. After confirmation of the fragmentation, the chromatin was diluted and incubated with NR2F1 antibody (10 μL) or isotype control IgG overnight. The antibody-enriched chromatin was then pelleted by magnet beads and purified with spin columns included in the kit. Fold enrichment of NR2F1 (relative to isotype control rabbit IgG) on the *Pth1r* promoter was determined by qPCR. The primer sequences are listed in **Supplemental Table 2**.

**Statistical analysis**

Clinical data were first imputed using 5 random forest iterations, each with 300 trees. Then, LASSO (least absolute shrinkage and selection operator) analyses were used to explore any associations between clinical data and cell-specific TGFBR2 expression in PCa patient bone metastasis tissues. Experimental data are presented as mean ± SEM. Longitudinal bone lesion data were analyzed via log transformed linear mixed-effects models with random slopes and intercepts. Bootstrap hypothesis testing with 1000 resampled data sets was used to test for differences in total lesion area between groups at specific time points. Normality assumptions were assessed visually via QQ-plots; no concerning deviations were detected. For bone lesion proportion analysis, we performed a two-sample proportions test on the proportion of the incidence of bone lesions in the two groups. The 95 percent confidence interval of the difference in the two groups is (0.2249110, 0.7473112), with a p-value of 0.0009424. For bone lesion area comparison in the *Tgfbr2^Col1CreERT^* KO and the *Tgfbr2^Flox^* mice tibia, since the bone lesion area distribution is not normal, we performed the Wilcoxon rank sum test with continuity correction and applied the Bonferroni-Holm multiple test correction. We found that the knockout group had significantly greater bone lesion area (*Mdn* = 3.4996) than the control group (*Mdn* = 0) at week 5, *W* = 137.5, *p* = 0.0414. For week 5.5, we found that the knockout group had significantly greater bone lesion area (*Mdn* = 5.2441) than the control group (*Mdn* = 0), *W* = 125.5, *p* = 0.0198. For further analysis only on tibiae with detectable lesions, we defined all tibiae with positive lesion area values at week 5.5 as “detectable” and analyzed the data on week 5 and week 5.5. Again, we performed the Wilcoxon rank sum test with continuity correction and applied the Bonferroni-Holm multiple test correction. We found the control group had greater bone lesion area at week 5 (*Mdn* = 7.4216) than the knockout group (*Mdn* = 3.8925), *W* = 89, *p* = 0.4261. For week 5.5, the control group had greater bone lesion area (*Mdn* = 9.9532) than the knockout group (*Mdn* = 5.5641), but the difference was not significant with *W* = 88, *p* = 0.4664.

Realtime qPCR results were analyzed via a linear mixed-effect model assuming unequal variances, on log2 transformed data with a random effect for each bio replicate and Benjamini-Hochberg false discovery rate adjustments for multiple testing. All the analyses were performed in R v 3.2.2. For all analyses, *p* < 0.05 was considered significant.

| **Supplementary Table 1**  Cell-specific TGFBR2 staining scores for the tissues from the bone metastatic tissue microarray. Provided as a separate file.  **Supplementary Table 2**  Primer sequences used in this article. | | |
| --- | --- | --- |
| **Primer ID** | **Application** | **Sequences (5'-3')** |
| mGapdh_F | cDNA expression | TCCTCAGTGTAGCCCAAGA |
| mGapdh_R | cDNA expression | GGAGAAACCTGCCAAGTATGA |
| mPth1r_F | cDNA expression | GCACACAGCAGCCAACATAA |
| mPth1r_R | cDNA expression | GGGGTAGAACTTTCCCGGTG |
| mNr2f1_F | cDNA expression | CTTACACATGCCGTGCCAAC |
| mNr2f1_R | cDNA expression | ATTCTTCCTCGCTGAACCGC |
| mNr2f1-p1F | ChIP | GGCTCTTCCCGACCCTAATG |
| mNr2f1-p2R | ChIP | CAGGCCTCGGGTTAAATTGC |
| mNr2f1-p2F | ChIP | GGGTGGACGGTAGCAAAGGA |
| mNr2f1-p2R | ChIP | AACCACACTGAGCCCTTCGG |
| mNr2f1-p1F | ChIP | GGCTCTTCCCGACCCTAATG |
| mNr2f1-p2R | ChIP | CAGGCCTCGGGTTAAATTGC |
| mAr-gRNA-1F | For Ar KD | CACCGggttacgccaaaggattgga |
| mAr-gRNA-1R | For Ar KD | AAACtccaatcctttggcgtaaccC |
| mAr-gRNA-2F | For Ar KD | CACCGacagacaagctcaaggatgg |
| mAr-gRNA-2R | For Ar KD | AAACccatccttgagcttgtctgtC |

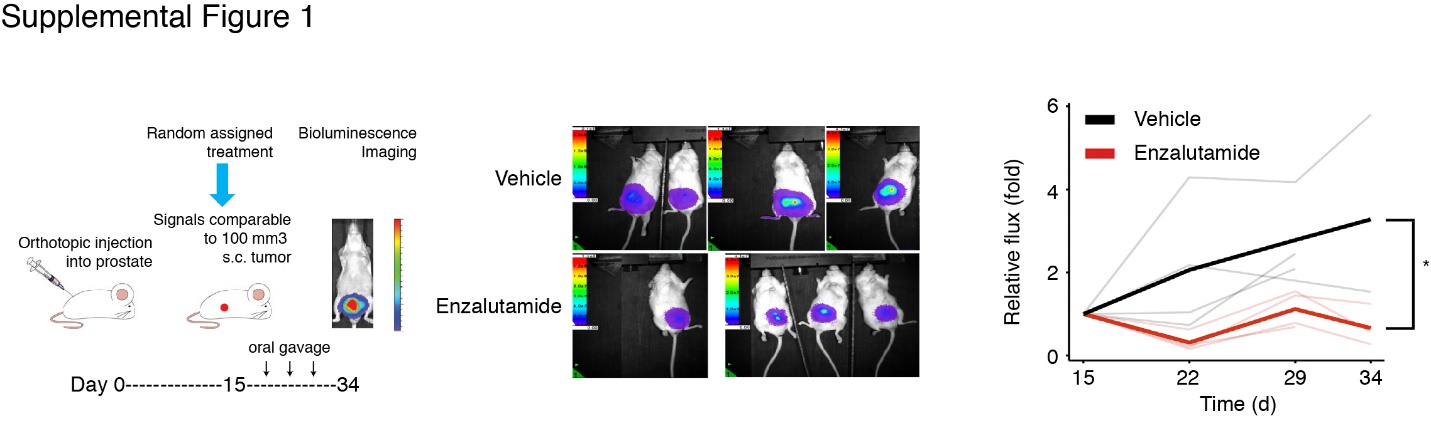
**Supplemental Figure 1**

Orthotopic tumors grown in the mouse prostate were imaged using bioluminescence once per week and analyzed using a linear mixed-effect model. *p* =0.023, *n* ≥ 4. The treatments were started on day 15 when the bioluminescent signals of orthotopic tumors were comparable to the signals from 100-mm^3^ subcutaneous tumors.

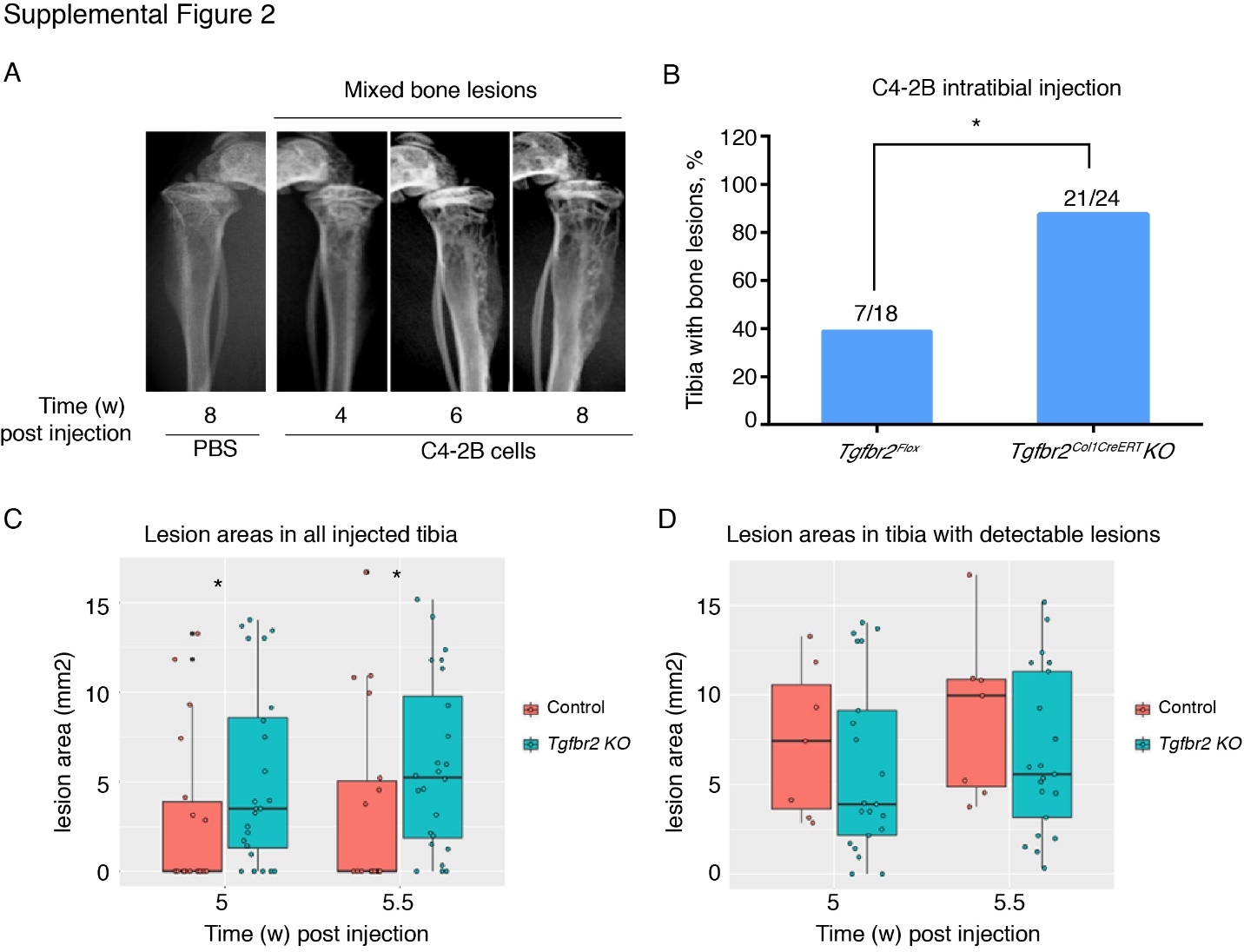

**Supplemental Figure 2**

**A)** Representative X-rays of mixed bone lesions of C4-2B cells after 4, 6, and 8 weeks post intratibial injection. **B)** The proportion of tibiae with detectable C4-2B cell-induced bone lesions in all the injected tibiae was significantly higher (*p*<0.001) in the *Tgfbr2^Col1CreERT^* KO group relative to the *Tgfbr2^Flox^* control group. The number of tibiae with detectable bone lesions identified by X-ray scanning of each group was counted 8 weeks after injection. The growth of C4-2B cells was confirmed by H&E staining (not shown). **C&D)** Bone lesion areas in all the injected tibiae **(C)** or only the tibiae with detectable lesions in control *Tgfbr2^Flox^* and *Tgfbr2^Col1CreERT^* KO mice. Lesion areas data collected at 5 weeks and 5.5 weeks post injection were plotted. For **C**, week 5, *p* = 0.0414; week 5.5, *p* = 0.0198. For **D**, week 5, *p* = 0.4261; week 5.5, *p* = 0.4664. A Wilcoxon rank sum test with continuity correction was performed. Details could be found in the Statistical Analysis section.

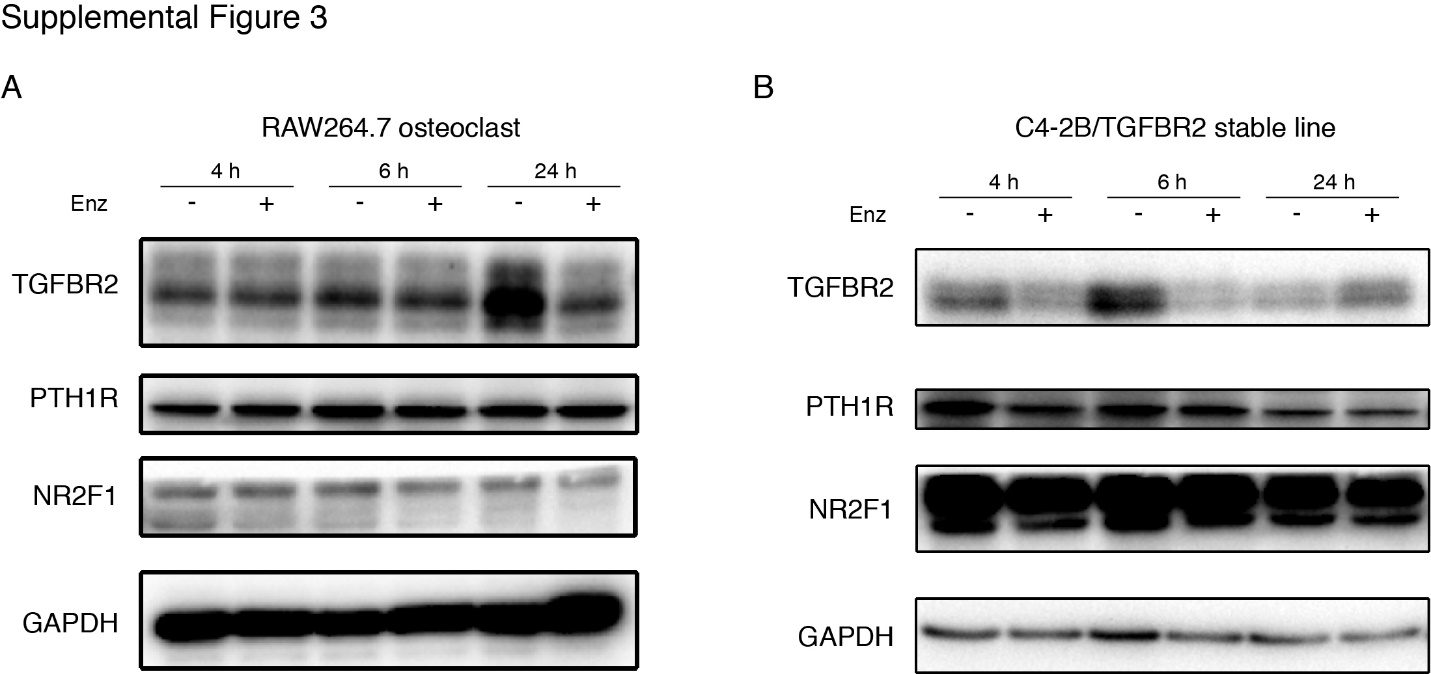

**Supplemental Figure 3**

Raw264.7 cells **(A)** and C4-2B/TGFBR2 cells **(B)** were treated with enzalutamide (Enz) or vehicle (DMSO) for 4 h, 6 h, or 24 h before harvesting for western blotting. Samples were blotted with TGFBR2, PTH1R, NR2F1, and GAPDH (loading control).
